# No evidence for transvection *in vivo* by a superenhancer:promoter pair integrated into identical open chromatin at the *Rosa26* locus

**DOI:** 10.1101/393363

**Authors:** Keiji Tanimoto, Hitomi Matsuzaki, Eiichi Okamura, Aki Ushiki, Akiyoshi Fukamizu, James Douglas Engel

## Abstract

Long-range associations between enhancers and their target gene promoters have been shown to play critical roles in executing genome function. Recent variations of chromosome capture technology have revealed a conprehensive view of intra- and inter-chromosomal contacts between specific genomic sites. The locus control region of the β-globin genes (β-LCR) is a super-enhancer that is capable of activating all of the β-like globin genes within the locus in *cis* through physical interaction by forming DNA loops. CTCF helps to mediate loop formation between LCR-HS5 and 3’HS1 in the human β-globin locus, in this way thought to contribute to the formation of a “chromatin hub”. The β-globin locus is also in close physical proximity to other erythrocyte-specific genes located long distances away on the same chromosome. In this case, erythrocyte-specific genes gather together at a shared “transcription factory” for co-transcription. Theoretically, enhancers could also activate target gene promoters on different chromosomes *in trans*, a phenomenon originally described as transvection in *Drosophilla*. Although close physical proximity has been reported for the β-LCR and the β-like globin genes when integrated at the mouse homologous loci in *trans*, their structural and functional interactions were found to be rare, possibly because a lack of suitable regulatory elements that might facilitate *trans* interactions. Therefore, we re-evaluated presumptive transvection-like enhancer-promoter communication by introducing CTCF binding sites and erythrocyte-specific transcription units into both LCR-enhancer and β-promoter alleles, each inserted into the mouse *ROSA26* locus on separate chromosomes. Following cross-mating of mice to place the two mutant loci at the identical chromosomal position and into active chromation in *trans*, their transcriptional output was evaluated. The results demonstrate that there was no significant functional association between the LCR and the β-globin gene *in trans* even in this idealized experimental context.

## Introduction

Gene expression is tightly regulated by DNA *cis* elements and their binding *trans*-factors, in which specific enhancer-promoter communications play a pivotal role. While genome-wide sequencing of the human and mouse genomes disclosed the number of genes to be more than 20,000, the number of enhancer elements is predicted to far exceed the number of genes [1]. Because accumulating evidence suggests that perturbation of enhancer function can be a major cause of pathogenesis in human diseases [2], it is of paramount importance to assign the activity of any individual enhancer to a specific target gene(s) in order to predict its function. Recent genome-wide interactome analyses revealed that enhancers can *physically* interact with genes over enormous distances, exceeding several hundreds of kilobase pairs in *cis*, or even with genes located on different chromosomes in *trans* [3], suggesting the presence of molecular mechanisms that allow specific enhancer-promoter interactions to take place over very long distances.

In the interphase nucleus, the genome adopts a higher-order chromatin architecture, in which transcription factors play important roles. Among those, CTCF, first identified as a transcriptional activator or repressor and subsequently, as an insulator, binds to two distinct genome regions to bring those two sites into close spatial proximity [4]. Ineractome analysis in ES cells revealed that the number of intra- or inter-chromosomal interactions mediated by CTCF was 1,480 and 336, respectively [5]. However, how frequently gene expression is reflected by changes in CTCF-mediated genome architecture is not well understood. On the other hand, it has been reported that genes with similar transcriptional specificity migrate into transcription factories in the nucleus that are rich in transcription factors engaged in the expression of those genes [6]. According to this mechanism, two distinct genome regions carrying genes with the same expression pattern should meet at the shared foci for co-transcription.

The human β-like globin genes are organized within a 70-kbp span on human chromosome 11, with the embryonic ε-globin gene located most 5’, followed by the two fetal γ-globin genes (Gγ and Aγ), while the adult δ- and β-globin genes are at the 3’ end of the locus (Fig. 1A). Expression of all the β-like globin genes in primitive, as well as in definitive erythroid cells, depends on the activity of the locus control region (LCR; [7, 8]), a super-enhancer element located 48 kbp 5’ to the transcription initiation site of the β-globin gene. The LCR consists of five DNaseI hypersensitive sites (HSSs), among which HS1 to 4 are constituent enhancers and rich in binding sites for transcription factors [9], while HS5 carries CTCF binding sites [10].

**Figure 1.**
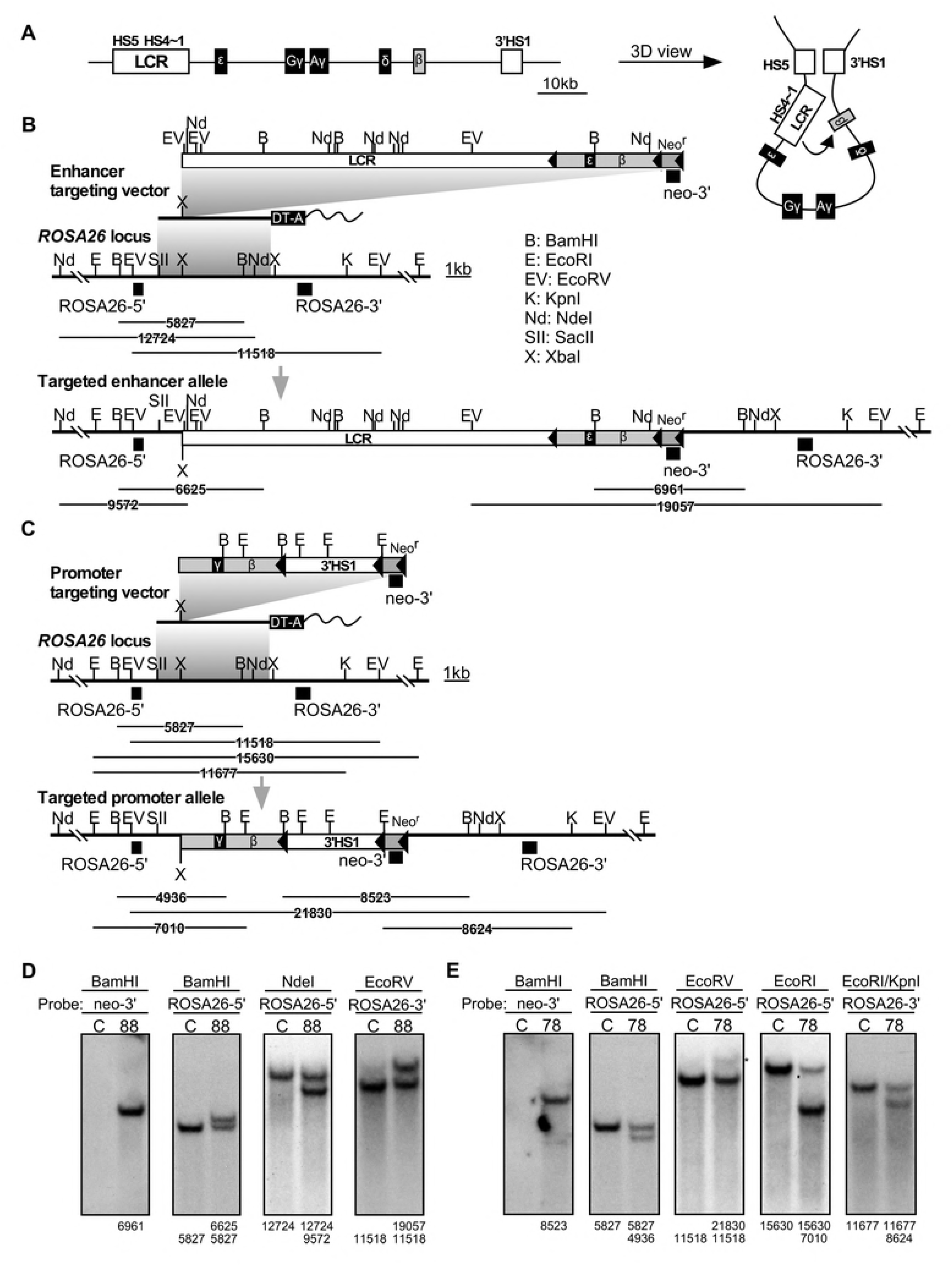
Generation of enhancer and promoter knock-in alleles in mice. **(A)** Structure of the human β-globin gene locus shown in 1D (left) and 3D (right) views. **(B)** The enhancer targeting vector carrying the human β-globin LCR and β-globin gene that is marked by an ε-globin sequence, wild-type *ROSA26* locus, and the correctly targeted enhancer knock-in locus are shown. In the targeting vector, neomycin resistance (Neo^r^) and diphtheria toxin (DT)-A genes are shown as striped and solid boxes, respectively. The solid triangles indicate the loxP sequences. Probes used for Southern blot analyses in (D) are shown as filled rectangles. Expected restriction fragments with their sizes are shown beneath the partial restriction enzyme maps. **(C)** The promoter targeting vector carrying the human β-globin gene (marked by an γ-globin sequence) and 3’HS1, wild-type *ROSA26* locus, and the correctly targeted promoter knock-in locus are shown. Probes used for Southern blot analyses in (E) are shown as filled rectangles. **(D and E)** Genomic DNA from ES clones was digested with restriction enzymes, separated on agarose gels, and Southern blots were hybridized to the probes. Asterisks denote nonspecific bands.

How the distal LCR enhancer activates β-globin gene expression has long been a subject of intense debate [11]. In 2002, RNA TRAP [12] and chromatin conformation capture (3-C; [13, 14]) assays elegantly revealed that the LCR and β-globin promoters were positioned in close proximity: these observations were consistent with a looping model, in which proteins bound to the LCR enhancer and to the gene promoters physically interact with the intervening DNA sequences looped out [15]. Erythroid specific transcription factors, such as GATA-1, NF-E2 and EKLF are essential for efficient globin genes transcription through binding to both the LCR and globin gene promoters. It is therefore presumed that they participate somehow in long-range enhancer-promoter interactions. In fact, both GATA-1 and NF-E2 are essential for LCR and βmaj-globin proximity in murine erythroid cells [16, 17], as well as for LCR and γ-globin proximity in human erythroid cells [18]. Similarly, EKLF is required for loop formation between the LCR and β-globin promoter sites, but not for LCR-HS5 and 3’HS1 sites [19]).

Interestingly, CTCF binding was found around the HSSs at both ends of the locus, *i.e*. LCR-HS5 and the 3’HS1 regions (Fig. 1A; [13]). Although 3C assays revealed proximal positioning of these sites in the nucleus, it was not confined to erythroid cells. In globin expressing cells, the LCR and 3 ‘HS1 regions are further located proximally to the actively expressed β-globin genes, which structure has been termed an active chromatin hub (Fig. 1A; [13, 20]). Therefore, transcriptional activation of the β-like globin genes is predicted to be a multi-step process, in which HS5-3’HS1 interaction may help to bring LCR enhancer sequences within close proximity of the β-globin promoter, thus facilitating their productive interaction (Fig. 1A).

It has been reported that non-genic LCR sequences are transcribed in erythroid cells [21, 22]. Therefore, LCR and β-globin gene may be co-transcribed in the same RNA polymerase II (PolII) factory, which then aids their physical association and transcriptional activation of the β-globin gene by the LCR enhancer. In accord with this notion, the β-globin gene locus on mouse chromosome 7 was found to colocalize with erythroid specific genes located 20 Mb away on the same chromosome in erythroid cell nuclei [6]. Furthermore, the murine β-globin gene locus colocalized with the *Slc4a1* (Chr. 11) and *Cd47* (Chr. 16) genes at a shared polII factory [23].

Because the LCR makes direct contact with its targets through loop formation, it can theoretically touch and activate such targets on separate chromosomes. Such *trans* chromosomal interaction for functional enhancer-promoter communicaton was dubbed transvection in *Drosophilla* [24, 25]. As mentioned above, recent development of biochemical technologies disclosed that interchromosomal *physical* interaction was common throughout the genome, not just in *Drosophilla* but also in mammals [26]. To test whether inter-chromosomal *functional* as well as physical association between the LCR enhancers and the β-like globin gene promoters take place, Noordermeer *et al*. knocked-in each sequence into the mouse genome separately on homologous chromosomes [27]. Because they found upregulation of endogenous murine β-like globin genes (~2-fold) on another chromosome, they concluded that the LCR must have some affinity for the β-globin promoter even in *trans*. However, neither frequent interchromosomal homologous chromosome interactions nor transvection-like activation of reporter genes in *trans* was observed,

In their experimental design, though, CTCF-assisted or co-transcriptionally mediated mechanisms were not considered. We therefore decided to re-evaluate transvection in mammals by incorporating well-characterized β-globin *cis* elements at the *ROSA26* locus.

Firstly, LCR-HS5 and 3’HS1 sequences were introduced into enhancer and promoter alleles, respectively, in expectation that CTCF factors bound at these sites might promote the formation of an interchromosomal bridge, which would in turn facilitate functional interactions between the LCR enhancer elements (HS4~1) and the β-globin promoter. In addition, β-globin transcription units were included in both alleles, anticipating that two alleles bearing the same promoter would likely migrate into same PolII factory in the nucleus. Even in this idealized experimental design, however, functional association between two alleles on separate chromosomes was not observed.

## Results

### Generation of enhancer and promoter knock-in alleles at the mouse *Rosa26* locus

A targeting vector for the enhancer allele (Fig. 1B, top) carried the LCR (HSs 1~5) and the β-globin gene sequences. The one for the promoter allele (Fig. 1C, top) carried the β-globin gene and the 3’HS1 sequences. To distinguish human β-globin gene transcripts expressed from each allele by PCR, portions of the β-globin gene were replaced either with corresponding segments of the ε- and γ-globin genes in the enhancer and promoter targeting vectors, respectively. In this experimental design, expression of both chimeric genes is under the control of a common human β-globin proximal promoter element (1.6 kb).

In the absence of a cis-linked LCR, the human β-globin gene locus becomes heterochromatinized (e.g. in the Hispanic thalassemia patient; [28, 29]) and the expression of all the β-like globin genes is reduced in transgenic mice [30, 31]. Therefore, a β-globin transgene without an LCR enhancer in *cis* on the promoter allele is expected to be heterochromatinized and to not be efficiently activated by the LCR on the enhancer allele in *trans*. We therefore chose the mouse *ROSA26* locus (on chromosome 6) for testing the transvection phenomenon, because this locus has a stable open chromatin structure in virtually all tissues [32].

To test for possible involvement, if any, of co-transcription of the β-globin genes in the enhancer and promoter alleles for transvection analysis, an ε-marked β-globin [β(ε)-globin] gene sequence was surrounded by loxP sites (floxed) in the enhancer allele so that it could be removed by conditional *in vivo* cre-loxP-mediated homologous recombination (Fig. 1B). To test for involvement of allelic proximity mediated by CTCF factors in the transvection experiment, the 3’HS1 sequence in the promoter allele was also floxed (Fig. 1C). Following homologous recombination with these targeting vectors in R1-ES cells, genomic DNA was prepared and correct recombination events were confirmed by Southern blot analyses using several combinations of restriction enzymes and specific probes (Fig. 1D and 1E).

### *In vivo* Cre-loxP recombination to derive daughter sublines

Following establishment of germ line modified mouse lines from the mutant ES cells by co-culture aggregation, the one carrying the enhancer knock-in allele (*LCR+β(ε)+Neo*^r^; Fig. 2A, top) was mated with cre-expressing TgM to remove either the “Neo^r^” or “Neo^r^ + *β(ε)*-globin gene” sequences, thereby generating either *“LCR+β(ε)’* or “*LCR*” alleles, respectively (Fig. 2A). Similarly, *“β(γ)+3’HS1* “ and “*β(γ)*” alleles were derived from lines carrying the promoter knock-in allele (*β(γ)+3’HS1+Neo^r^*) by removing “Neo^r^” or “Neo^r^ +3’HS1” sequences, respectively (Fig. 2B). Correct cre-loxP recombination events were confirmed by Southern blotting (Fig. 2C), as well as PCR analyses of tail genomic DNAs (Fig. 2D).

**Figure 2.**
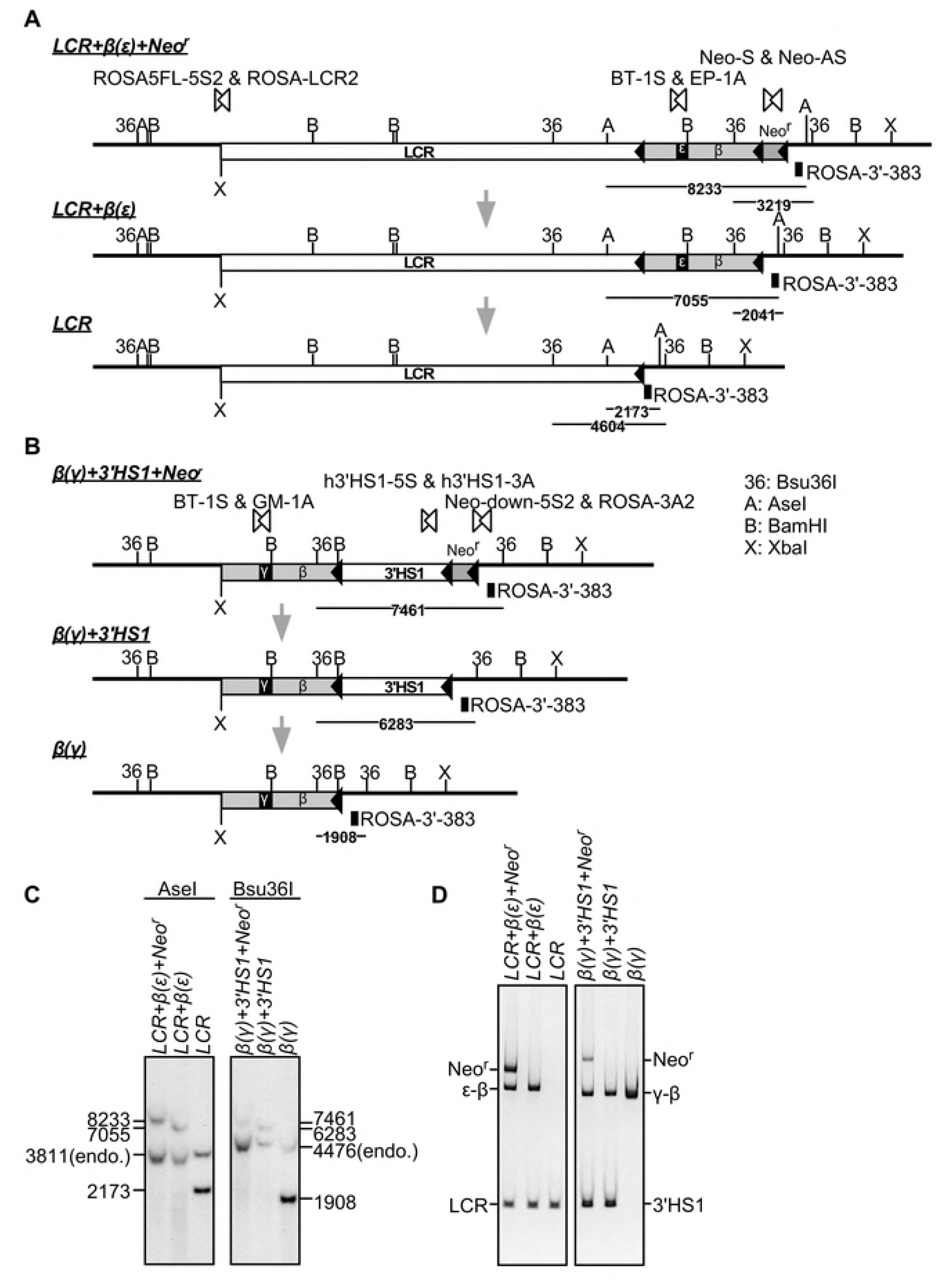
Derivation of enhancer/promoter-allele variants by *in vivo* Cre-loxP recombination. **(A)** Enhancer knock-in mouse bearing the *LCR+β(ε)+Neo^r^* locus was mated with Cre-TgM to induce *in utero*, partial *cre-loxP* recombination, which resulted in selective excision of either Neo^r^ or Neo^r^+β(ε)-globin sequences to generate *LCR+β(ε)* or *LCR* alleles, respectively. A, AseI; B, *BamHI;* 36, Bsu36I. **(B)** Similarly, *β(y)+3?Sl* or *β(γ)* alleles were derived from the promoter knock-in mouse bearing the *β(γ)+3’HS1+Neo^r^* locus by deletion of the Neo^r^ or Neo^r^+3’HS1 sequences, respectively. **(C)** Successful cre-loxP recombination was confirmed by Southern blot analysis. Tail genomic DNA of mutant mice were digested with AseI (enhancer knock-in series) or Bsu36I (promoter knock-in series), separated on agarose gels, and Southern blots were hybridized to the ROSA-3’-383 probe shown in panels A and B. **(D)** Each allele was discriminated by multiplex PCR analyses of tail genomic DNA from mutant mice. The LCR, ε-β and Neo^r^ sequences in the enhancer knock-in alleles were amplified by PCR primers shown by paired open arrowheads in panel A. The γ-β, 3’HS1 and Neo^r^ sequences in the promoter knock-in alleles were amplified by PCR primers shown by paired open arrowheads in panel B.

### Evaluation of enhancer activity *in vivo*

To analyze chimeric β-globin gene expression in the knock-in mutant alleles, adult mice were made anemic, nucleated erythroid cells were collected from their spleens and recovered RNAs were reverse-transcribed. Expression of hybrid β(γ)- and β(ε)-globin genes was analyzed using a primer set comon to both chimeric gene sequences. Addition of 3 ‘HS1 to the β(γ)-globin gene increased its expression level by only 1.6-fold *in vivo* (Fig. 3A), which was less prominent when compared with its effect in YAC TgM [33]. In human β-globin YAC TgM, deletion of 3’HS1 sequences attenuated adult β-globin gene expression by more than 10-fold, possibly because this sequence plays additional roles in higher order chromatin organization at the human β-globin locus in the 150-kb YAC TgM [34].

**Figure 3.**
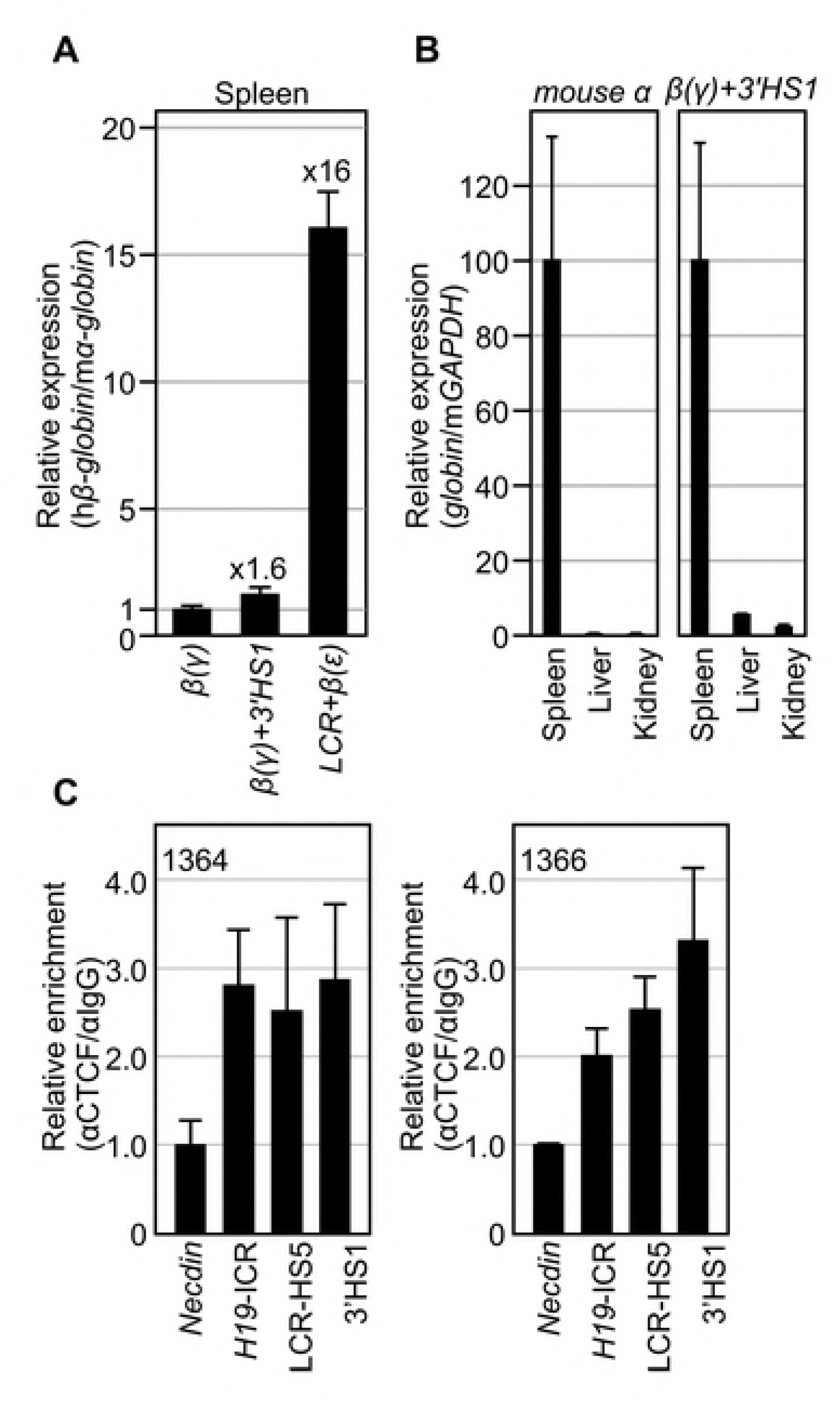
Expression of human β-globin genes in knock-in mice. **(A)** Total RNA was prepared from spleens of 1-month-old anemic mice (N=4 for each genotype). Expression levels of the human β(γ)- or β(ε)-globin genes (analyzed by common primer set targeted at β-globin sequence) and endogenous mouse α (mα)-globin gene were analyzed by semi-quantitative RT-PCR. The ratio of hβ/mα-globin genes was calculated (the expression value of the β(γ)-globin was set at 1). **(B)** Total RNA was prepared from spleens, livers and kidneys of 1-month-old anemic mice (N=4 for each tissues). Expression of mα-globin, β(γ)-globin and endogenous mouse (m)GAPDH genes was analyzed by semi-quantitative RT-PCR. The expression levels of mα- (left panel) or β(γ)-globin (right) genes, both compared to that of mGAPDH gene were calculated (the expression values in spleen samples were set at 100). **(C and D)** ChIP was conducted for CTCF in the spleen cells of anemic animals bearing both *LCR+β(ε)* and *β(γ)+3’HS1* alleles. The *Necdin* gene and the *H19* ICR sequences were analyzed as negative and positive controls, respectively. Quantitative PCR was repeated at least three times for each sample. Fold enrichment of CTCF relative to IgG control (average values with S.D.) was calculated and graphically depicted (average value of negative controls was set at 1.0).

The expression level of the β(ε)-globin gene linked to the LCR in *cis* was 16-fold higher than that of the β(γ)-globin gene in isolation (Fig. 3A). This magnitude seemed much less significant than when LCR is deleted from the whole locus in endogenous [29] or transgenic environments [35]. It is generally accepted that the LCR not only potentiates promoter activity as an activator but also opens chromatin [29]. While this latter activity is a part of the “enhancer” function in the context of the native β-globin locus, the *Rosa26* locus is in an open chromatin configuration by its nature and therefore, the observed 16-fold activation may represent a promoter potentiation function of the LCR alone.

### Evaluation of read-through transcription from the *Rosa26* promoter

Because the enhancer and promoter constructs were integrated at the ubiquitously expressed *Rosa26* gene locus, some portion of the chimeric β-globin gene transcription could be driven by the *ROSA26* gene promoter. We therefore quantified how much read-through transcription from the *Rosa26* gene promoter might contribute to expression of the β-globin gene sequence (Fig. 3B). Total RNA was extracted from the spleen, liver and kidney of anemic adult mice and the expression levels of β(γ)-globin+3’HS1 gene and the mouse α-globin gene, relative to the GAPDH gene expression, were determined by qRT-PCR. The mouse α-globin gene was preferentially expressed only in the spleen, as expected (Fig. 3B, left). In contrast, while β(γ)-globin+3’HS1 was highly expressed in the spleen, its low level expression was also observed in the liver and kidney. Since α-globin gene expression was barely detected in these non-hematopoietic tissues (in adults), we concluded that contamination of erythroid cells in these tissues was negligible. Therefore, low level β(γ)-globin+3’HS1 gene expression in the liver and kidney (and probably in the spleen) was under the control of the *Rosa26* promoter. In other words, it appears that at least 90% of β(γ)-globin+3’HS1 gene transcription in the spleen initiates from the β-globin gene promoter.

### Confirmation of CTCF binding to the LCR-HS5 and 3’HS1 regions

To test for CTCF binding to well-established CTCF binding sites in LCR-HS5 and 3’HS1 of the human β-globin locus in mutant animals, ChIP analyses were conducted using chromatin prepared from anemic spleen (erythroid) cells (Fig. 3C). PCR primers for *H19* ICR and *necdin* [4] loci were included as positive and negative controls for CTCF binding, respectively.

As expected, CTCF enrichment was observed at the LCR-HS5 and 3 ‘HS1 regions, confirming its binding to these sites *in vivo*.

### Cross-mating to derive animals carrying distinct pairs of enhancer and promoter alleles

To generate four distinct combinations of enhancer- and promoter-knock-in alleles (Fig. 4A), animals carrying heterozygous enhancer alleles (*LCR^+/−^* or *LCR+β(ε)-globin^+/−^)* and those carrying homozygous promoter alleles (*β(γ)-globin^+/+^* or *β(γ)-globin+3 ‘HS1^+/+^*) were mated. Theoretically, in the next generation, half of the litters carry both enhancer and promoter alleles in *trans* and the other half carry promoter allele only (Fig. 4A). Derivation of animals bearing the expected genotypes was confirmed by allele-specific PCR (Fig. 4B; results for animals with promoter allele alone is not shown) and Southern blot analyses (data not shown).

**Figure 4.**
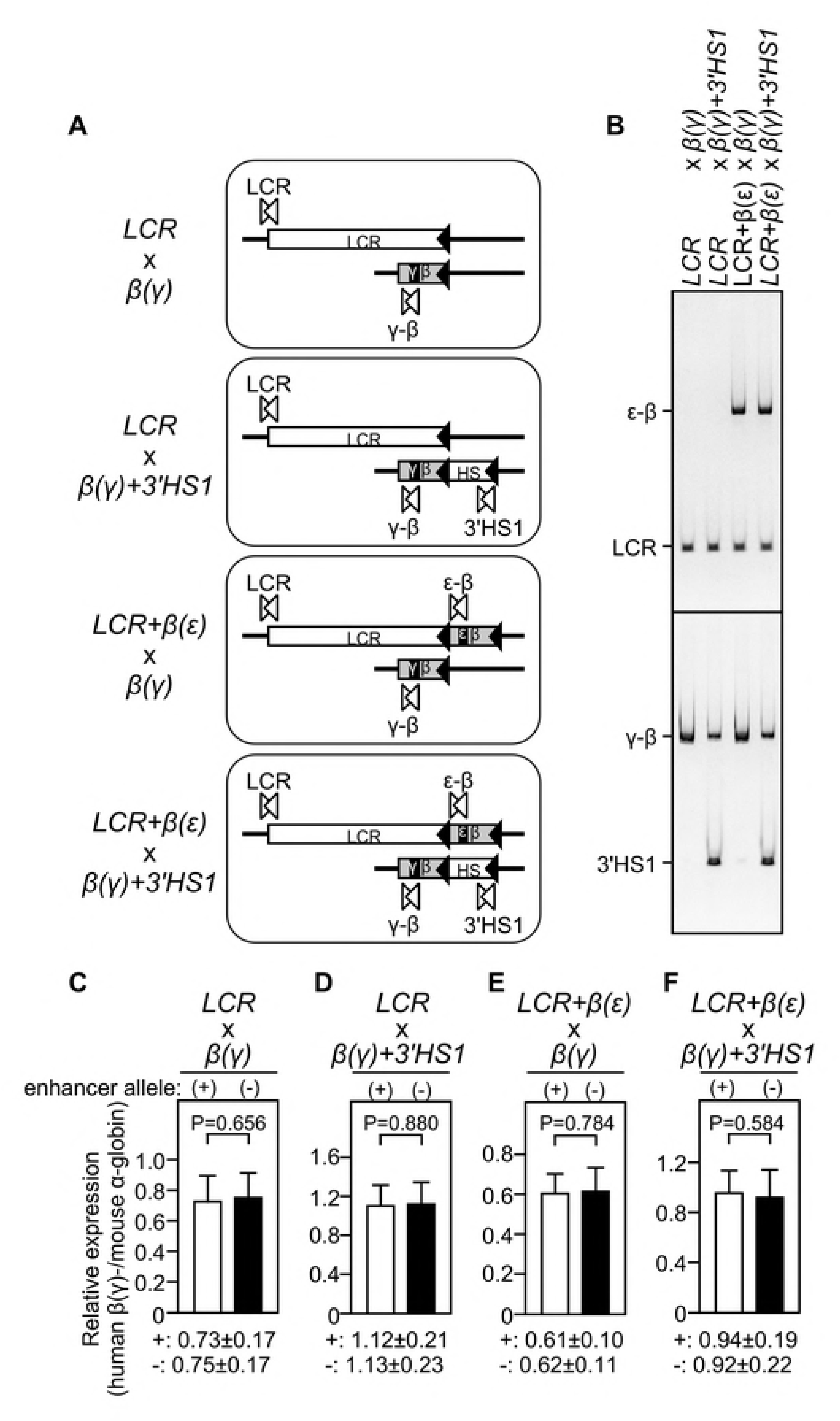
Expression of β(γ)-globin genes in knock-in mice. **(A)** Schematic representation of four different combinations of enhancer and promoter knock-in alleles to test for enhancer-promoter interaction *in trans*. **(B)** Mouse genotypes shown in (A) were confirmed by multiplex PCR analyses of tail genomic DNA of mutant mice. The ε-β, LCR, γ-β and 3’HS1 sequences were amplified by PCR primers shown by paired open arrowheads in panel A. **(C-F)** Total RNA was prepared from spleens of 1-month-old anemic mice. Numbers analyzed for each genotype are shown in the Supplemental Figure 1. Expression of β(γ)-globin and endogenous mα-globin genes was analyzed by semi-quantitative RT-PCR. The ratio of hβ(γ)-globin / mα-globin genes was calculated and average value with S.D. was graphically depicted for each genotype group (Although values are arbitrary, they can be quantitatively compared between the panels).

### The test for functional interchromosomal enhancer-promoter interactions

The expression of the hybrid β(γ)-globin gene was compared in two animal groups carrying either the promoter allele alone or both enhancer and promoter alleles in *trans* (Fig. 4C-F and Supplemental Figure 1). Anemic spleens were collected from ~1 month old animals and the expression levels of human β-globin and mouse α-globin genes were determined by qRT-PCR.

Human β(γ)-globin gene expression normalized to that of the mouse α-globin gene in the two groups (with or without the LCR *in trans*) did not differ significantly, providing no evidence for trans-activation (Fig. 4C and Supple. Fig. 1A). Even when compared within single litters, no significant difference was observed between two groups (data not shown). This result was consistent with a report by Noordermeer *et al* [27], in which the LCR and γ-globin gene were individually integrated at a gene-dense site on mouse chromosome 8 and tested for inter-chromosomal interaction.

Next, the β(γ)-globin gene with an attached 3’HS1 sequence was used as a reporter and the experiment was repeated (Fig. 4D and Supple. Fig. 1B). Because the LCR-HS5 and 3’HS1 sequences were bound by CTCF (Fig. 3C), it was possible that the LCR and 3’HS1 come into close proximity, which then facilitates trans-activation of β(γ)-globin gene by the LCR. Even in this idealized experimental setting, however, no significant activation was observed.

Then, the combination of LCR+β(ε)-globin and β(γ)-globin genes as enhancer and promoter alleles, respectively, was tested (Fig. 4E and Supple. Fig. 1C). Because transcription units in these two alleles share the same transcriptional regulatory sequences (*i.e*. the β-globin proximal promoter), it was possible that two alleles would migrate into a shared transcription factory, which would then lead to β(γ)-promoter activation by the LCR enhancer in *trans*, caused by the close proximity of the two alleles. As shown in Fig. 4E, however, no sign of trans-activation was detected.

Finally, the combination of the LCR+β(ε)-globin and β(γ)-globin+3’HS1 alleles was investigated (Fig. 4F and Supple. Fig. 1D). Although in some of the litters, statistically significant difference in the expression levels in between promoter alone and enhancer+promoter alleles was observed (Supple. Fig. 1D), this significant but subtle difference dissapeared when the sample number increased.

## Discussion

Long-range chromatin interactions can occur over long distance between regions on the same chromosome in *cis* or on distinct chromosomes in *trans*. Because chromatin clustering events are dynamic, consequential formation of higher-order genome organization is believed to play an important role in regulating gene expression [36, 37]. In *Drosophila*, a pair of homologous chromosomes can physically interact to allow productive enhancer-promoter communication in *trans*, a phenomenon referred to as transvection [24, 25]. In the case of the *yellow* locus, for example, an enhancer of one copy of a gene (that lacks promoter activity) regulates the expression of the paired copy of the gene (lacking enhancer activity) in *trans*. More recently, many inter-chromosomal interactions were identified using 3C-based biochemical strategies in mammalian cells [38], and growing evidence suggests that transvection-like phenomena may be generally possible for eukaryotic enhancer-promoter communication.

Several mechanisms have been proposed for establishing preferential interactions between genomic loci, including enhancer-promoter communication via specific sets of transcription factors, co-migration of gene loci with similar transcriptional profiles into shared PolII foci, or reflection of structural constraints mediated by architectural proteins such as CTCF/cohesin. These mechanisms may also account for inter-chromosomal chromatin interactions, such as those observed in the interferon-γ gene [39], the mouse *Hoxb1* gene [40], and the housekeeping *Rad23a* gene [41]. Among such examples, the *H19* imprinting control region *(H19* ICR) on mouse chromosome 7 has been shown to interact with regions on several different chromosomes in *trans*, including the maternally inherited insulin-like growth factor 2 receptor *(Igf2r)* on chromosome 17, to suppress its transcription [42]. Because deletion of the *H19* ICR affected the expression of interacting genes on different chromosomes, interchromosomal interaction was suggested play a functional role in their transcriptional regulation.

To gain insight into possible inter-chromosomal gene regulatory mechanisms, Noordermeer *et al*. employed an LCR super-enhancer and human β-like globin promoter as regulatory elements to test for functional consequences *(i.e*. gene transcription) of placing them separately at corresponding *cis* locations on homologous chromosomes in mice [27] since these regulatory elements represent one of the most robust and most thoroughly examined enhancer-promoter pairs capable of interacting with each other over extremely long distances [6]. Despite their clear affinity in the native chromatin configuration, they did not show significant inter-chromosomal interaction when integrated into the *Rad23a* gene locus [41]. Our observations reported here are consistent with their results (*LCR* = *β(γ)* in Fig. 4C).

Because enhancer-promoter looping in the β-globin gene locus is dependent on erythroid-specific transcription factors, such as EKLF and GATA-1, it has been anticipated that long-range chromatin loops were formed only in the cells where target genes are active [13]. In fact, in the EKLF knock-out mice, significantly reduced chromatin loop formation between the LCR and β-globin gene was accompanying diminished β-globin gene transcription [19]. Recent observations in mice, however, suggest that chromatin connectivity is not necessarily coupled to transcriptional events. For example, sonic hedgehog (Shh) gene transcription is regulated by a distal enhancer and they form a chromatin loop. This loop, however seems to be preset and detectable even when *Shh* is not transcribed [43]. In addition, among several interactions between regulatory sequences within the *HoxD* locus, some are present even in the absence of target gene activation [44, 45]. In the β-globin gene locus, formation of an active chromatin hub leading to erythroid-specific gene activation turned out to be a multistep process; *e*. the chromatin loop between the LCR-HS5 and 3’HS1 was preformed in erythroid progenitor cells in a CTCF-dependent fashion prior to globin gene transcription [13]. Importantly, this developmentally early structure was not affected by EKLF ablation in the previously cited knock-out experiment [19]. It can therefore be assumed that, upon erythroid cell maturation, this prestructure (outer loop) may facilitate subsequent gene activation that accompanies inner loop formation between an enhancer and promoter, only when cell type-specific transcription factors are present.

By using a cell-based system, in which nuclear levels of GATA-1 can be experimentally manipulated, it was shown that β-globin gene activity, as well as the contact between the LCR and the gene required nuclear GATA-1 [16]. The knockdown of LDB1, a cofactor of GATA-1, also impaired both β-globin gene transcriptional activity and LCR-β-globin loop formation, presumably by affecting GATA-1 function [46]. Interestingly, when LDB1 was tethered to the β-globin promoter region via an artificial zinc-finger motif, chromatin looping and β-globin gene activity was partially restored, even in the absence of GATA1 [47]. Furthermore, forced loop formation between the LCR and the γ-globin promoter in adult erythroid cells caused ectopic upregulation of γ-globin transcription [48]. These instances demonstrate that engineered formation of a chromatin loop alone between an enhancer and promoter was sufficient to activate gene expression. In other words, certain forms of chromatin interactions may play an instructive role in gene expression and set the permissive condition for actual gene activation.

It has been reported that some fraction of CTCF binding sites were located at enhancer and promoter regions and involved in their physical associations [49]. To facilitate enhancer-promoter interactions in *trans*, we therefore placed binding sites for CTCF in both enhancer and promoter alleles in our experiment. As mentioned above, it has been reported that the LCR-HS5 and the 3’HS1 are in close proximity in both globin-expressing and non-expressing erythroid cells, suggesting that they have physical affinity independent of β-globin gene expression. However, we failed to observe upregulation of β(γ)-globin gene expression even in this idealized condition (Fig. 4D & F). While 3C analysis of our human β-globin YAC TgM clearly identified a chromatin loop between the LCR and the β-globin promoter in *cis*, any sign of *trans*-interaction was not validated by our current work (data not shown). Recently, Hi-C analysis of the human genome demonstrated that chromatin loops were often established between two opposed CTCF sites in opposite motif directions [50]. Furthermore, as for the establishment of CTCF/cohesin-mediated loops, an “extrusion model” has been proposed [51], in which a loop-extruding molecule *(i.e*. cohesin) forms progressively larger loops but are stalled at CTCF-bound sites *(i.e*. boundary elements). If this is the singular mechanism that allows CTCF-mediated interaction between distinct genomic sites, inter-chromosomal association via CTCF/cohesin machinery cannot be possible. However, because a massive number of CTCF-associated inter-chromosomal interactions have been reported [5], intervening chromatin should not absolutely be required for the formation of chromatin interactions. Nevertheless, addition of CTCF sites on both enhancer and promoter knock-in alleles did not augment their functional *trans* association in our experiments.

The majority of protein coding genes are transcribed by RNA polymerase II (PolII) and this process is also engaged in long-range associations between gene loci [52]. It is conceivable that coregulated genes (and enhancers) with common temporal and spatial specificities may migrate to preassembled PolII factories for transcription and therefore their loci can be in close proximity, even when they are separated by long distance in *cis* on the same chromosome or in *trans* on chromosome pairs. Such a situation has been reported for genes on mouse chromosome 7, which are involved in globin synthesis and regulation. They are transcriptionally active in erythroid cells and their loci co-localized at shared PolII foci with the active β-globin genes. [6, 23]. Therefore, preferential long-range interactions between distinct gene loci can be generated by genes being transcribed. In other words, two genomic elements with spatial proximity in the nuclear space would have a chance to interact with each other. Although it has been reported that the LCR is transcribed in erythroid cells [21], the enhancer knock-in allele employed by Noordermeer *et al*. was unlikely to be transcribed efficiently because of a lack of the ERV-9 LTR sequence [22]. In addition, although the *Rad23a* is a housekeeping gene, it is not known if the knock-in alleles in that study were efficiently transcribed. We therefore introduced β-globin transcriptional regulatory units both in enhancer and promoter knock-in alleles at the *ROSA26* locus (*LCR+β(ε)* x *β(γ)* or β(γ)+3’HS1 in Fig. 4A). Despite their significant expression in erythroid cells (Fig. 3A & B), however, we did not observe increased reporter gene expression when compared to that in the absence of a paired enhancer allele in *trans* (Fig. 4E & F).

In the examples of the *Shh* and the *HoxD* loci, higher order pre-structures have been proposed to facilitate future promoter–enhancer contacts over very long distances [43, 44]. *Cis*-interaction between the LCR-HS5 and 3’HS1 found at the endogenous β-globin locus (outer loop) may not be sufficiently stable to facilitate LCR-β promoter interaction (inner loop) when they are situated in *trans*. It is also possible that the function of the outer loop may simply be to isolate a given locus from its surrounding chromatin environment [4, 13, 53]). Further functional evaluation of the higher order chromatin structure is apparently required to ask if “form always follows function”.

## Materials and Methods

### Targeting vectors

Floxed neomycin resistance gene (flNeo^r^) cassete was released from pMC1neopA_5’/3’-loxP [54] by *XbaI/NheI* digestion and inserted into *XbaI* site of pROSA26-1 (generous gift of Dr. Philippe Soriano; nucleotide position at 180,029 in AC155722; RP24-204J8) to generate pRO SA26/MC 1neopA_5 ‘/3’-loxP(-).

5’-upstream portion of human β-globin gene (nucleotides 60,577-60,882 in HUMHBB; U01317.1; GenBank) was PCR-amplified by using a set of oligonucleotides: ICI-02-5S; 5’-GG<u>GGTACC</u> <u>TCTAGATCT</u>CTATTTATTTAGCA-3’ (artificial <u>*Kpn*I</u> and <u>*Xba*I</u>, and endogenous <u>*Bgl*II</u> [at 60,557] sites are underlined) and ICI-02-3A; 5’-GGTCAGCGTAGGGTCTCAGT-3’. Following *Kpn*I and *Apa*I (at 60,882) digestion, this fragment and 3’-downstream portion of the gene (*Apa*I-*Xba*I fragment; nucleotides 60,882-65,426) were linked and cloned into *Kpn*I/*Xba*I sites of pBluescriptII KS(+). *Bam*HI site (at 60,676) of this plasmid was then eliminated to generate pβ-globin_K-X-ΔB for facilitating cloning procedure.

Portions of ε- and Aγ-globin gene sequences were PCR-amplified by using following two sets of oligonucleotides: ICI-04-5S;

5’-GGCACCATGGTGCATTTTACTGCT-3’ (artificial *Nco*I site underlined) and ICI-03-3A; 5’-TCAGGATCCACATGCAGCTT-3’ (<u>*Bam*HI</u>) or ICI-03-5S;

5’-CACACACTCGCTTCTGGAAC-3’ and ICI-03-3A, respectively. Following *Nco*I (partial) and BamHI digestion, ε (nucleotides 19,539-19,959) and Aγ (nucleotides 39,465-39,885) gene sequences were replaced with corresponding portion of β-globin gene (nucleotides 62,185-62,613) in pβ-globin_K-X-ΔB to generate pmβ/ε and pmβ/γ, respectively.

[Promoter targeting vector]

Following two oligonucleotides were annealed, phosphorylated and inserted into *Nde*I site (at 65,287 in HUMHBB) of the β-globin gene in pmβ/γ to generate pmβ/γ_loxP(-): BTLX-5S; 5’-TATC<u>GGATCCT</u>*ATAACTTCGTATAATGTATGCTATACGAAGTTA*TAGA-3’ and BTLX-3A; 5’-TATCTA *TAACTTCGTA TAGCA TACA TTA TACGAAGTTA* <u>TAGGATCC</u>GA-3’. In each oligo, loxP sequences are italisized and BamHI sites underlined. The *Xba*I fragment, carrying β-globin promoter, γ sequence-marked β-globin coding region, and a loxP site, was released from pmβ/γ_loxP(-) and introduced into XbaI site of pROSA26/MC1neopA_5’/3’-loxP(-) to generate pR26/loxP-Neo/mβ/γ.

Human β-globin 3’HS1 sequence (4,194 bp, *SmaI-HindIII*) was subcloned from BAC clone, RP11-1205H24 (nucleotides 35,934-40,127 in AC129505; GenBank). Upon conversion of *HindIII* site to SmaI site, the fragment was introduced into *Sma*I site of pR26/loxP-Neo/mβ/γ to generate pR26/loxP-Neo/mβ/γ/3’HS1 (Promoter targeting vector).

[Enhancer targeting vector]

Following two oligonucleotides were annealed, phosphorylated and inserted into BglII site (at 60,557 in HUMHBB) of the β-globin gene in pmβ/ε to generate pmβ/ε_loxP(-): ICI-08-5S; 5’-GATC<u>GGCGCGCC</u>*ATAACTTCGTATAATGTATGCTATACGAAGTTAT-3’* and ICI-08-3A; 5’-GATC*ATAACTTCGTATAGCATACATTA*TACGAAGTTATGGCGCGCC-3’. In each oligo, loxP sequences are italisized and *Asc*I sites underlined. The XbaI fragment, carrying a loxP site, β-globin promoter, and ε sequence-marked β-globin coding region, was released from pmβ/ε_loxP(-) and introduced into XbaI site of pROSA26/MC1neopA_5’/3’-loxP(-) to generate pR26/loxP-Neo/mβ/ε.

For facilitating cloning procedure, following two oligonucleotides were annealed, phosphorylated and inserted into *KpnI/SacI* site of pBluescriptII KS(+): ICI-01-5S; 5’- GGCGCGCCGGTACCTATGCGGCCGCGGCGCGCCAGCT-3’ and ICI-01-3A; 5’-GGCGCGCCGCGGCCGCATAGGTACCGGCGCGCCGTAC-3’, which resulted in *AscI-KpnI-NotI-[SacII]-AscI* sites formation. Human β-globin LCR sequences (17,590 bp, EcoRI-XbaI) were recovered from pRS/LCR [55] as *KpnI-NotI* (both in the multicloning sites) fragment and inserted into *KpnI-NotI* site of above plasmid to generate pBS/LCR. Upon eliminating SacII site (parenthesized site in the above double-stranded oligo) from pBS/LCR, 3’ downstream region of HS1 was accidentally deleted and the final LCR size cloned was (17,198 bp, nucleotides 95,131-77,934 in AC104389; Ensemble). Finally, LCR sequences were recovered from pBS/LCR as AscI fragment and introduced into AscI site of pR26/loxP-Neo/mβ/ε to generate pR26/LCR/loxP-Neo/mβ/ε (Enhancer targeting vector).

### Gene targeting in ES cells and generation of mutant mice

Target vectors were linearized by SacII digestion. R1-ES cells were grown on embryonic fibroblast feeder cells. Following electroporation (Bio-Rad GenePulser Xcell [0.4 mm gap] at setting of 250 V and 500 microfarads) of cells (1.0 = 10^7^ cells) with a linearized targeting vector (20 μg), cells were selected in 0.4 mg/ml G418. Homologous recombination in ES cells was first screened by PCR and then confirmed by Southern blotting with several combinations of restriction enzymes and probes shown below.

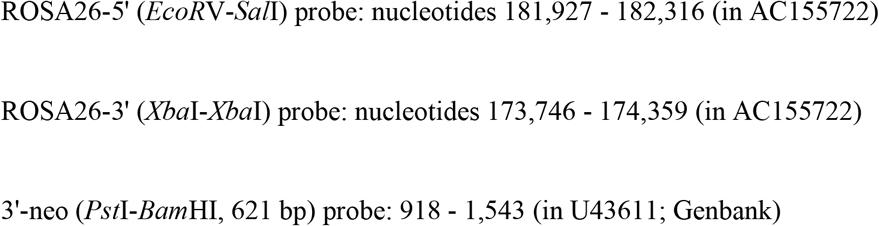

Chimeric mice were generated by a coculture method using eight-cell embryos from CD1 mice (ICR, Charles River Laboratories), bred with CD1 females, and germ line transmission of the mutant allele was determined by PCR and Southern blot analyses.

TgM ubiquitously expressing cre recombinase were mated with knock-in mice to partially or completely execute Cre-loxP recombination, which was confirmed by PCR and Southern blot analyses of tail DNA of offsprings.

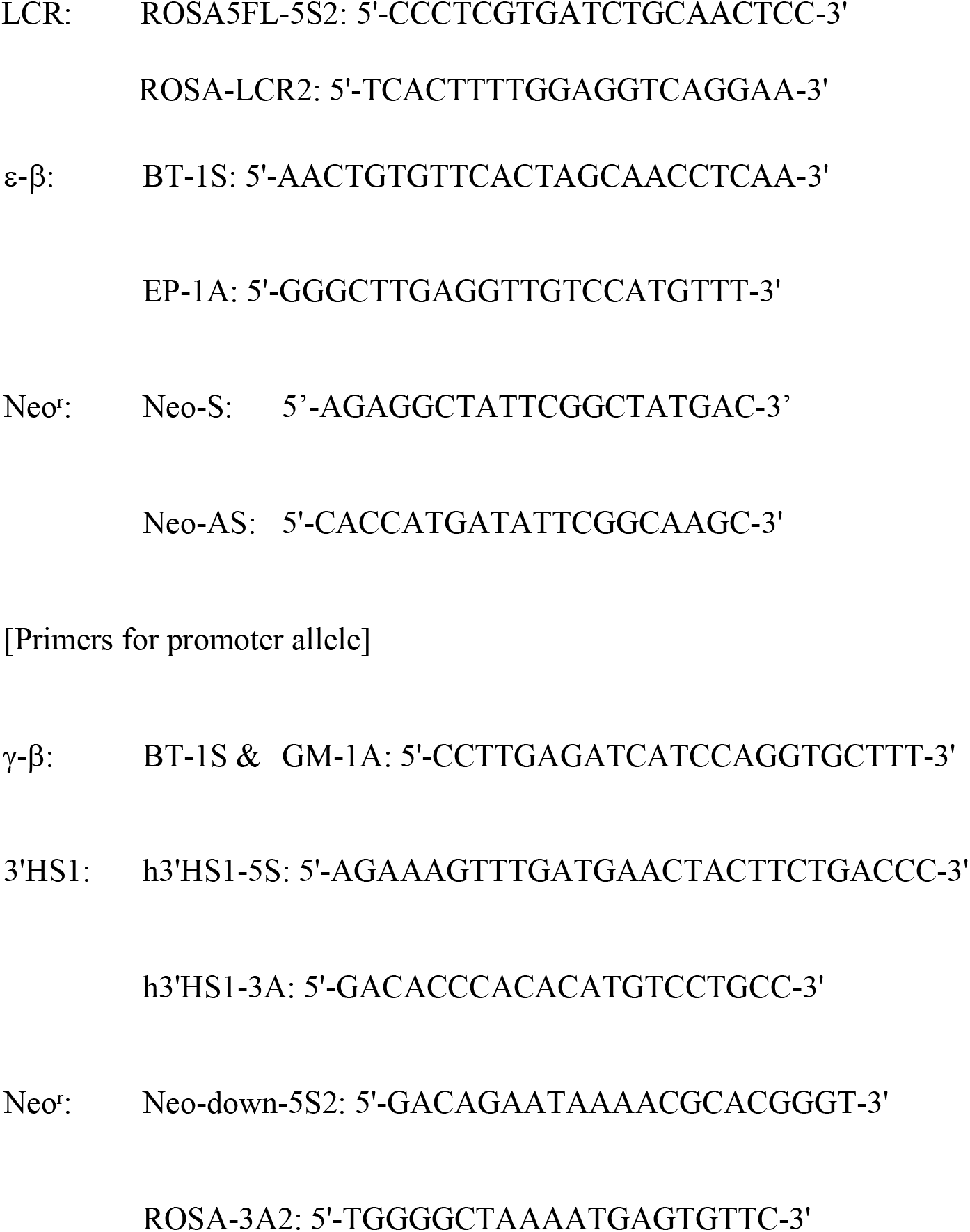

### Animal procedures

Mice were housed in a pathogen-free barrier facility in a 12-hour light/12-hour dark cycle, and fed standard rodent chow. Adult mice were sacrificed by cervical dislocation and the organs were immediately removed and flash-frozen in liquid nitrogen.

Animal experiments were performed in a humane manner under approval from the Institutional Animal Experiment Committee of the University of Tsukuba. Experiments were performed in accordance with the Regulation of Animal Experiments of the University of Tsukuba and the Fundamental Guidelines for Proper Conduct of Animal Experiments and Related Activities in Academic Research Institutions under the jurisdiction of the Ministry of Education, Culture, Sports, Science and Technology of Japan.

### Expression analysis

Total RNA was extracted from phenyl hydrazine-induced anemic adult spleens (1 to 2 months old) by ISOGEN (Nippon Gene) and converted to cDNA using ReverTra Ace qPCR RT Master Mix with gDNA Remover (TOYOBO). One-fortieth of the reaction mixture was subjected to quantitative PCR amplification using the KOD SYBR qPCR Mix (Toyobo) and thermal Cycler Dice (TaKaRa Bio) with the following parameters: 95°C for 5s and 60°C for 30s, 40 cycles.

The PCR primer sets used for human β(γ)- or β(ε)-globin genes amplification were GM-1S2 and BT-3A3 (126-bp amplicon) or BT-1S3 and EP-3A (152-bp), respectively. Primer set common to both β(γ)- and β(ε)-globin genes amplification in Fig. 3A was BT-4S1 and BT-4A1. The primer sets used for internal control gene expression analyses were MAgI-I and II (mouse α-globin gene) or mGAPDH-5S and 3A (mouse *gapdh* gene).

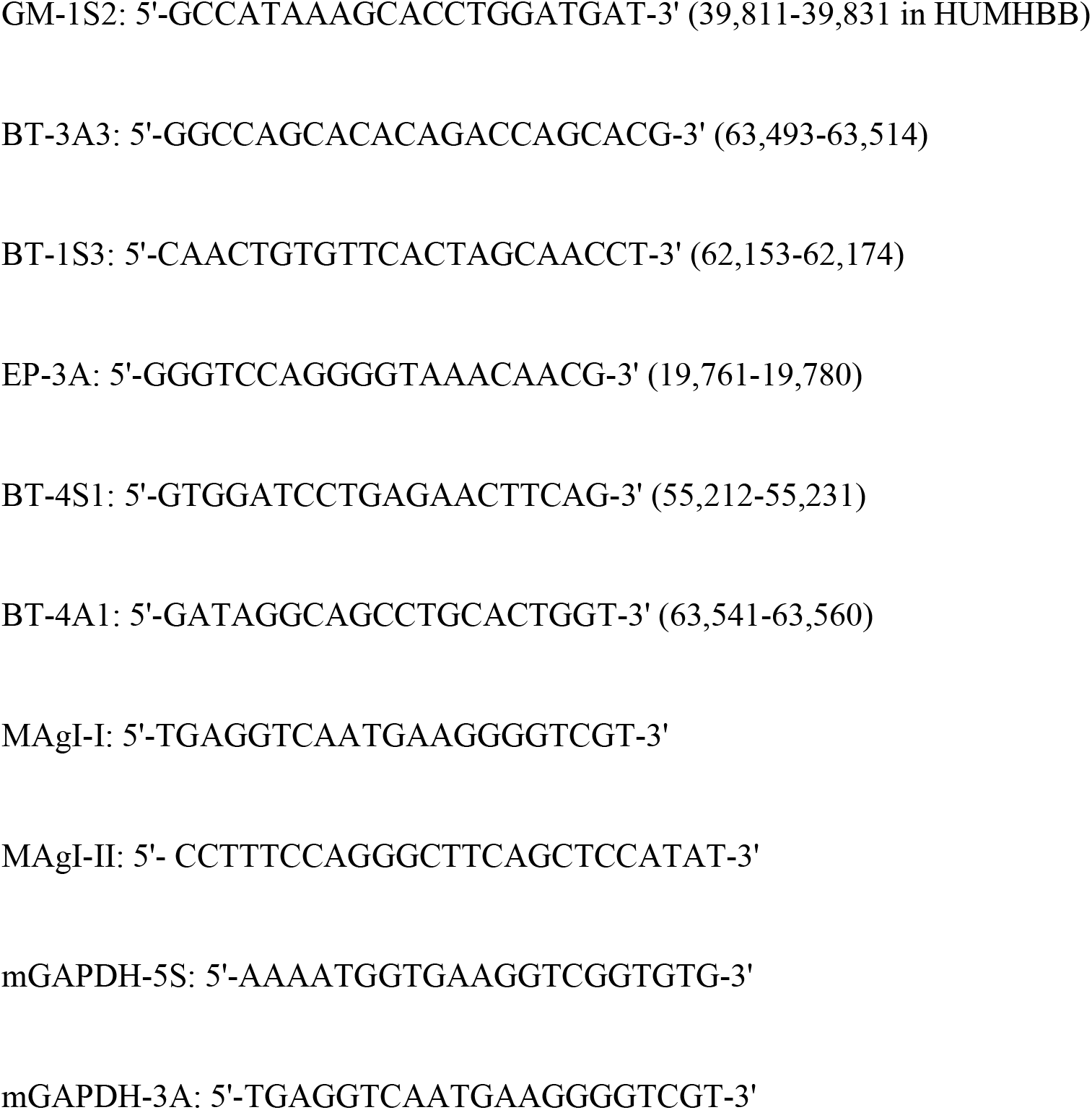

### ChIP analysis

The animals (2 to 4 months old) bearing both the *LCR+β(ε)* and *β(γ)+3’HS1* knock-in alleles in *trans* were made anemic and nucleated erythroid cells were collected from their spleens. Following fixation with 1% formaldehyde for 10 min at room temperature. Nuclei (2 = 10^7^ cells) were digested with 12.5 units/ml of micrococcal nuclease at 37°C for 20 min. The chromatin was incubated with anti-CTCF antibody (D31H2; Cell Signaling Technology) or purified rabbit IgG (Invitrogen) overnight at 4°C and was precipitated with preblocked Dynabeads protein G magnetic beads (Life Technologies, Carlsbad, CA). Immunoprecipitated materials were then washed and reverse cross-linked. DNA was purified with the QIAquick PCR purification kit (Qiagen, Venlo, The Netherlands) and subjected to qPCR analysis. The endogenous *H19* ICR and *Necdin* sequences were analyzed as positive and negative controls, respectively [56]. LCR-HS5 and 3’HS1 primer sets are as follows:

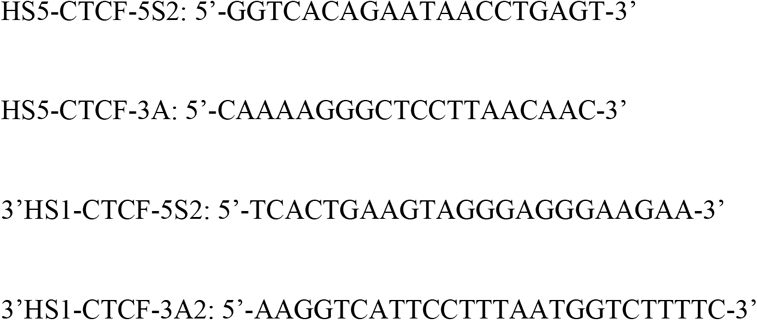

## References

1. Consortium EP. An integrated encyclopedia of DNA elements in the human genome. Nature. 2012;489(7414):57–74. doi: 10.1038/nature11247. PubMed PMID: 22955616; PubMed Central PMCID: PMCPMC3439153.

2. Smith E, Shilatifard A. Enhancer biology and enhanceropathies. Nat Struct Mol Biol. 2014;21(3):210–9. doi: 10.1038/nsmb.2784. PubMed PMID: 24599251.

3. Chakalova L, Debrand E, Mitchell JA, Osborne CS, Fraser P. Replication and transcription: shaping the landscape of the genome. Nat Rev Genet. 2005;6(9):669–77.Epub 2005/08/12. doi: 10.1038/nrg1673. PubMed PMID: 16094312.

4. Splinter E, Heath H, Kooren J, Palstra RJ, Klous P, Grosveld F, et al. CTCF mediates long-range chromatin looping and local histone modification in the beta-globin locus. Genes Dev. 2006;20(17):2349–54. Epub 2006/09/05. doi: 10.1101/gad.399506. PubMed PMID: 16951251; PubMed Central PMCID: PMCPMC1560409.

5. Handoko L, Xu H, Li G, Ngan CY, Chew E, Schnapp M, et al. CTCF-mediated functional chromatin interactome in pluripotent cells. Nat Genet. 2011;43(7):630–8. Epub 2011/06/21. doi: 10.1038/ng.857. PubMed PMID: 21685913; PubMed Central PMCID: PMCPMC3436933.

6. Osborne CS, Chakalova L, Brown KE, Carter D, Horton A, Debrand E, et al. Active genes dynamically colocalize to shared sites of ongoing transcription. Nat Genet. 2004;36(10):1065–71. Epub 2004/09/14. doi: 10.1038/ng1423. PubMed PMID: 15361872.

7. Epner E, Reik A, Cimbora D, Telling A, Bender MA, Fiering S, et al. The beta-globin LCR is not necessary for an open chromatin structure or developmentally regulated transcription of the native mouse beta-globin locus. Mol Cell. 1998;2(4):447–55. Epub 1998/11/11. PubMed PMID: 9809066.

8. Li Q, Peterson KR, Fang X, Stamatoyannopoulos G. Locus control regions. Blood. 2002;100(9):3077–86. doi: 10.1182/blood-2002-04-1104. PubMed PMID: 12384402; PubMed Central PMCID: PMCPMC2811695.

9. Hnisz D, Shrinivas K, Young RA, Chakraborty AK, Sharp PA. A Phase Separation Model for Transcriptional Control. Cell. 2017;169(1):13–23. doi: 10.1016/j.cell.2017.02.007. PubMed PMID: 28340338; PubMed Central PMCID: PMCPMC5432200.

10. Tanimoto K, Sugiura A, Omori A, Felsenfeld G, Engel JD, Fukamizu A. Human beta-globin locus control region HS5 contains CTCF-and developmental stage-dependent enhancer-blocking activity in erythroid cells. Mol Cell Biol. 2003;23(24):8946–52. PubMed PMID: 14645507; PubMed Central PMCID: PMCPMC309639.

11. Bulger M, Groudine M. Looping versus linking: toward a model for long-distance gene activation. Genes Dev. 1999;13(19):2465–77. PubMed PMID: 10521391.

12. Carter D, Chakalova L, Osborne CS, Dai YF, Fraser P. Long-range chromatin regulatory interactions in vivo. Nat Genet. 2002;32(4):623–6. Epub 2002/11/12. doi: 10.1038/ng1051. PubMed PMID: 12426570.

13. Palstra RJ, Tolhuis B, Splinter E, Nijmeijer R, Grosveld F, de Laat W. The beta-globin nuclear compartment in development and erythroid differentiation. Nat Genet. 2003;35(2):190–4. Epub 2003/10/01. doi: 10.1038/ng1244. PubMed PMID: 14517543.

14. Tolhuis B, Palstra RJ, Splinter E, Grosveld F, de Laat W. Looping and interaction between hypersensitive sites in the active beta-globin locus. Mol Cell. 2002;10(6):1453–65. Epub 2002/12/31. PubMed PMID: 12504019.

15. Choi OR, Engel JD. Developmental regulation of beta-globin gene switching. Cell. 1988;55(1):17–26. PubMed PMID: 3167976.

16. Vakoc CR, Letting DL, Gheldof N, Sawado T, Bender MA, Groudine M, et al. Proximity among distant regulatory elements at the beta-globin locus requires GATA-1 and FOG-1. Mol Cell. 2005;17(3):453–62. Epub 2005/02/08. doi: 10.1016/j.molcel.2004.12.028. PubMed PMID: 15694345.

17. Du MJ, Lv X, Hao DL, Zhao GW, Wu XS, Wu F, et al. MafK/NF-E2 p18 is required for beta-globin genes activation by mediating the proximity of LCR and active beta-globin genes in MEL cell line. Int J Biochem Cell Biol. 2008;40(8):1481–93. Epub 2008/03/01. doi: 10.1016/j.biocel.2007.11.004. PubMed PMID: 18308612.

18. Woon Kim Y, Kim S, Geun Kim C, Kim A. The distinctive roles of erythroid specific activator GATA-1 and NF-E2 in transcription of the human fetal gamma-globin genes. Nucleic Acids Res. 2011;39(16):6944–55. Epub 2011/05/26. doi: 10.1093/nar/gkr253. PubMed PMID: 21609963; PubMed Central PMCID: PMCPMC3167640.

19. Drissen R, Palstra RJ, Gillemans N, Splinter E, Grosveld F, Philipsen S, et al. The active spatial organization of the beta-globin locus requires the transcription factor EKLF. Genes Dev. 2004;18(20):2485–90. Epub 2004/10/19. doi: 10.1101/gad.317004. PubMed PMID: 15489291; PubMed Central PMCID: PMCPMC529536.

20. Patrinos GP, de Krom M, de Boer E, Langeveld A, Imam AM, Strouboulis J, et al. Multiple interactions between regulatory regions are required to stabilize an active chromatin hub. Genes Dev. 2004;18(12):1495–509. Epub 2004/06/17. doi: 10.1101/gad.289704. PubMed PMID: 15198986; PubMed Central PMCID: PMCPMC423198.

21. Routledge SJ, Proudfoot NJ. Definition of transcriptional promoters in the human beta globin locus control region. J Mol Biol. 2002;323(4):601–11. PubMed PMID: 12419253.

22. Long Q, Bengra C, Li C, Kutlar F, Tuan D. A long terminal repeat of the human endogenous retrovirus ERV-9 is located in the 5’ boundary area of the human beta-globin locus control region. Genomics. 1998;54(3):542–55. doi: 10.1006/geno.1998.5608.PubMed PMID: 9878258.

23. Schoenfelder S, Sexton T, Chakalova L, Cope NF, Horton A, Andrews S, et al. Preferential associations between co-regulated genes reveal a transcriptional interactome in erythroid cells. Nat Genet. 2010;42(1):53–61. Epub 2009/12/17. doi: 10.1038/ng.496. PubMed PMID: 20010836; PubMed Central PMCID: PMCPMC3237402.

24. Tartof KD, Henikoff S. Trans-sensing effects from Drosophila to humans. Cell. 1991;65(2):201–3. PubMed PMID: 2015622.

25. Duncan IW. Transvection effects in Drosophila. Annu Rev Genet. 2002;36:521–56. doi: 10.1146/annurev.genet.36.060402.100441. PubMed PMID: 12429702.

26. Belton JM, McCord RP, Gibcus JH, Naumova N, Zhan Y, Dekker J. Hi-C: a comprehensive technique to capture the conformation of genomes. Methods. 2012;58(3):268–76. doi: 10.1016/j.ymeth.2012.05.001. PubMed PMID: 22652625; PubMed Central PMCID: PMCPMC3874846.

27. Noordermeer D, de Wit E, Klous P, van de Werken H, Simonis M, Lopez-Jones M, et al. Variegated gene expression caused by cell-specific long-range DNA interactions. Nat Cell Biol. 2011;13(8):944–51. Epub 2011/06/28. doi: 10.1038/ncb2278. PubMed PMID: 21706023; PubMed Central PMCID: PMCPMC3151580.

28. Driscoll MC, Dobkin CS, Alter BP. Gamma delta beta-thalassemia due to a de novo mutation deleting the 5’ beta-globin gene activation-region hypersensitive sites. Proc Natl Acad Sci U S A. 1989;86(19):7470–4. PubMed PMID: 2798417; PubMed Central PMCID: PMCPMC298086.

29. Forrester WC, Epner E, Driscoll MC, Enver T, Brice M, Papayannopoulou T, et al. A deletion of the human beta-globin locus activation region causes a major alteration in chromatin structure and replication across the entire beta-globin locus. Genes Dev. 1990;4(10):1637–49. PubMed PMID: 2249769.

30. Magram J, Chada K, Costantini F. Developmental regulation of a cloned adult beta-globin gene in transgenic mice. Nature. 1985;315(6017):338–40. PubMed PMID: 3858676.

31. Townes TM, Lingrel JB, Chen HY, Brinster RL, Palmiter RD. Erythroid-specific expression of human beta-globin genes in transgenic mice. EMBO J. 1985;4(7):1715–23. PubMed PMID: 2992937; PubMed Central PMCID: PMCPMC554408.

32. Soriano P. Generalized lacZ expression with the ROSA26 Cre reporter strain. Nat Genet. 1999;21(1):70–1. doi: 10.1038/5007. PubMed PMID: 9916792.

33. Liu Q, Bungert J, Engel JD. Mutation of gene-proximal regulatory elements disrupts human epsilon-,gamma-,and beta-globin expression in yeast artificial chromosome transgenic mice. Proc Natl Acad Sci U S A. 1997;94(1):169–74. PubMed PMID: 8990180; PubMed Central PMCID: PMCPMC19271.

34. Fang X, Xiang P, Yin W, Stamatoyannopoulos G, Li Q. Cooperativeness of the higher chromatin structure of the beta-globin locus revealed by the deletion mutations of DNase I hypersensitive site 3 of the LCR. J Mol Biol. 2007;365(1):31–7. doi: 10.1016/j.jmb.2006.09.072. PubMed PMID: 17056066; PubMed Central PMCID: PMCPMC2826273.

35. Grosveld F, van Assendelft GB, Greaves DR, Kollias G. Position-independent, high-level expression of the human beta-globin gene in transgenic mice. Cell. 1987;51(6):975–85. PubMed PMID: 3690667.

36. Dekker J. Gene regulation in the third dimension. Science. 2008;319(5871):1793–4. doi: 10.1126/science.1152850. PubMed PMID: 18369139; PubMed Central PMCID: PMCPMC2666883.

37. Bonev B, Cavalli G. Organization and function of the 3D genome. Nat Rev Genet. 2016;17(12):772. doi: 10.1038/nrg.2016.147. PubMed PMID: 28704353.

38. Dekker J, Misteli T. Long-Range Chromatin Interactions. Cold Spring Harb Perspect Biol. 2015;7(10):a019356. doi: 10.1101/cshperspect.a019356. PubMed PMID: 26430217; PubMed Central PMCID: PMCPMC4588061.

39. Spilianakis CG, Lalioti MD, Town T, Lee GR, Flavell RA. Interchromosomal associations between alternatively expressed loci. Nature. 2005;435(7042):637–45. doi: 10.1038/nature03574. PubMed PMID: 15880101.

40. Wurtele H, Chartrand P. Genome-wide scanning of HoxB1-associated loci in mouse ES cells using an open-ended Chromosome Conformation Capture methodology. Chromosome Res. 2006;14(5):477–95. doi: 10.1007/s10577-006-1075-0. PubMed PMID: 16823611.

41. Simonis M, Klous P, Splinter E, Moshkin Y, Willemsen R, de Wit E, et al. Nuclear organization of active and inactive chromatin domains uncovered by chromosome conformation capture-on-chip (4C). Nat Genet. 2006;38(11):1348–54. doi: 10.1038/ng1896. PubMed PMID: 17033623.

42. Zhao Z, Tavoosidana G, Sjolinder M, Gondor A, Mariano P, Wang S, et al. Circular chromosome conformation capture (4C) uncovers extensive networks of epigenetically regulated intra-and interchromosomal interactions. Nat Genet. 2006;38(11):1341–7. doi: 10.1038/ng1891. PubMed PMID: 17033624.

43. Amano T, Sagai T, Tanabe H, Mizushina Y, Nakazawa H, Shiroishi T. Chromosomal dynamics at the Shh locus: limb bud-specific differential regulation of competence and active transcription. Dev Cell. 2009;16(1):47–57. doi: 10.1016/j.devcel.2008.11.011. PubMed PMID: 19097946.

44. Montavon T, Soshnikova N, Mascrez B, Joye E, Thevenet L, Splinter E, et al. A regulatory archipelago controls Hox genes transcription in digits. Cell. 2011;147(5):1132–45. doi: 10.1016/j.cell.2011.10.023. PubMed PMID: 22118467.

45. Andrey G, Montavon T, Mascrez B, Gonzalez F, Noordermeer D, Leleu M, et al. A switch between topological domains underlies HoxD genes collinearity in mouse limbs. Science. 2013;340(6137):1234167. doi: 10.1126/science.1234167. PubMed PMID: 23744951.

46. Song SH, Hou C, Dean A. A positive role for NLI/Ldb1 in long-range beta-globin locus control region function. Mol Cell. 2007;28(5):810–22. doi: 10.1016/j.molcel.2007.09.025. PubMed PMID: 18082606; PubMed Central PMCID: PMCPMC2195932.

47. Deng W, Lee J, Wang H, Miller J, Reik A, Gregory PD, et al. Controlling long-range genomic interactions at a native locus by targeted tethering of a looping factor. Cell. 2012;149(6):1233–44. doi: 10.1016/j.cell.2012.03.051. PubMed PMID: 22682246;PubMed Central PMCID: PMCPMC3372860.

48. Deng W, Rupon JW, Krivega I, Breda L, Motta I, Jahn KS, et al. Reactivation of developmentally silenced globin genes by forced chromatin looping. Cell. 2014;158(4):849–60. doi: 10.1016/j.cell.2014.05.050. PubMed PMID: 25126789; PubMed Central PMCID: PMCPMC4134511.

49. Shen Y, Yue F, McCleary DF, Ye Z, Edsall L, Kuan S, et al. A map of the cis-regulatory sequences in the mouse genome. Nature. 2012;488(7409):116–20. doi: 10.1038/nature11243. PubMed PMID: 22763441; PubMed Central PMCID:PMCPMC4041622.

50. Rao SS, Huntley MH, Durand NC, Stamenova EK, Bochkov ID, Robinson JT, et al. A 3D map of the human genome at kilobase resolution reveals principles of chromatin looping. Cell. 2014;159(7):1665–80. Epub 2014/12/17. doi: 10.1016/j.cell.2014.11.021. PubMed PMID: 25497547; PubMed Central PMCID: PMCPMC5635824.

51. Fudenberg G, Imakaev M, Lu C, Goloborodko A, Abdennur N, Mirny LA. Formation of Chromosomal Domains by Loop Extrusion. Cell Rep. 2016;15(9):2038–49. doi: 10.1016/j.celrep.2016.04.085. PubMed PMID: 27210764; PubMed Central PMCID: PMCPMC4889513.

52. Mitchell JA, Fraser P. Transcription factories are nuclear subcompartments that remain in the absence of transcription. Genes Dev. 2008;22(1):20–5. doi: 10.1101/gad.454008. PubMed PMID: 18172162; PubMed Central PMCID: PMCPMC2151011.

53. Oti M, Falck J, Huynen MA, Zhou H. CTCF-mediated chromatin loops enclose inducible gene regulatory domains. BMC Genomics. 2016;17:252. Epub 2016/03/24. doi: 10.1186/s12864-016-2516-6. PubMed PMID: 27004515; PubMed Central PMCID: PMCPMC4804521.

54. Matsuzaki H, Okamura E, Takahashi T, Ushiki A, Nakamura T, Nakano T, et al. De novo DNA methylation through the 5’-segment of the H19 ICR maintains its imprint during early embryogenesis. Development. 2015;142(22):3833–44. doi: 10.1242/dev.126003. PubMed PMID: 26417043.

55. Leach KM, Nightingale K, Igarashi K, Levings PP, Engel JD, Becker PB, et al. Reconstitution of human beta-globin locus control region hypersensitive sites in the absence of chromatin assembly. Mol Cell Biol. 2001;21(8):2629–40. doi: 10.1128/MCB.21.8.2629-2640.2001. PubMed PMID: 11283243; PubMed Central PMCID: PMCPMC86894.

56. Okamura E, Matsuzaki H, Sakaguchi R, Takahashi T, Fukamizu A, Tanimoto K. The H19 imprinting control region mediates preimplantation imprinted methylation of nearby sequences in yeast artificial chromosome transgenic mice. Mol Cell Biol. 2013;33(4):858–71. doi: 10.1128/MCB.01003-12. PubMed PMID: 23230275; PubMed Central PMCID: PMCPMC3571351.

